# Factors associated with misdiagnosis of hospitalised patients in general hospitals of Central Uganda

**DOI:** 10.1101/2022.09.16.508252

**Authors:** Simon Peter Katongole, Patricia Akweongo, Robert Anguyo DDMO, Daniel Evans Kasozi, Augustine Adoma Afari

## Abstract

Misdiagnosis of inpatients is a major public health issue whose scope and causes are unknown in Sub-Saharan African countries. The purpose of this cross-sectional study, which was conducted in five hospitals in central Uganda, was to identify the factors associated with inpatient misdiagnosis in general hospitals in Central Uganda. Records of 2,431 patients admitted between July 1st, 2019 and June 30th, 2020 were specifically reviewed to obtain data on variables thought to be associated with misdiagnosis. The admission diagnosis assigned at the emergency or outpatient department was compared to the discharge diagnosis assigned immediately after the patient’s admission, with any difference considered a misdiagnosis. The disease, patient, health system, and environmental factors associated with misdiagnosis were identified using multivariable logistic regression analysis.

Misdiagnosis was found in the records of 223/2431 (9.2%) of the admitted patients. A patient admitted to Nakaseke hospital [aOR=1.95, 95% CI=1.17-3.25, p=0.01], being admitted at night [aOR=3, 95% CI=1.81-5.02, p0.01], male patient [aOR=1.89, 95% CI=1.35-2.64, p0.01], patient’s age groups 10-19 [AOR=2.3, 95% CI=2.3-9.25, p0.01]; 20-29 [AOR=8.15, 95% CI=4.18-15.89], p<0.01; 30-39; and 40-49;; AOR=8.12, 95% CI=3.99-16.54, p<0.01; AOR=7.88, 95% CI=3.71-16.73, p<0.01; and AOR=12.14, 95% CI=6.41-23.01, p<0.0]. Misdiagnosis was also associated with multimorbidity (aOR=4.71, 95% CI=1.91-11.65, p0.01) and patients treated for uncommon diseases (aOR=2.57, 95% CI=1.28-5.18, p0.01). Patients without underlying diseases [aOR=0.63; 95% CI=0.43-0.91, p=0.015] and those who were not referred [aOR=0.51; 95% CI=0.31-0.86, p=0] .011] were less likely to be related to misdiagnosis.

To improve diagnostic accuracy, hospitals should reorganize patient admission processes, conducted targeted training, develop policy or guidelines targeting factors predisposing to misdiagnosis, and the adopt a diagnostic error prevention culture.

## Background

Over 5-15% of patients in hospitals worldwide are misdiagnosed, resulting in poor quality care and poor health outcomes ^1^. However, available research in low-income countries provides insufficient evidence to assist practitioners and policymakers in fully comprehending the burden that misdiagnosis imposes on clinical practice and health-care delivery ^2,3^. While misdiagnosis is generally underreported in Sub-Saharan Africa, it has been reported for some diseases such as HIV to range from 0.1% to 6.6% ^4^. In another study in Uganda5, 30.6% of patients with HIV-associated lymphoma were misdiagnosed as having tuberculosis ^5^. In Malawi, a misdiagnosis of cancer in 34 patients initially diagnosed with tuberculosis underscores the magnitude of the problem^6^. as many as 62% of febrile illnesses in rural Uganda were diagnosed as malaria when malaria was not present ^7^. However, because the majority of these studies are disease-specific, generalizing the findings can be difficult.

When a diagnostic error occurs, most people are interested in who was involved and what happened next. Less emphasis is placed on the factors that could be associated with the occurrence of the error ^8^. However, patient harm caused by misdiagnosis would be avoided more effectively if the factors associated with misdiagnosis were established by empirical evidence obtained from the departments, organizations, or health care systems where the misdiagnosis is occurring. This study thus identifies and explains the mechanisms by which admitted patients are misdiagnosed in Ugandan hospitals, in order to inform processes aimed at improving patient diagnosis in Ugandan general hospitals and related healthcare systems in low-income and Sub-Saharan African contexts.

## Methods

### Study design

This was a cross-sectional study that involved retrospective review of patients records that had been admitted between July 1^st^ 2019 to June 30^th^ 2020.

#### Study sites

The research was carried out at Mityana, Gombe, Nakaseke, Kiboga, and Kayunga general hospitals in central Uganda. In Uganda, a general hospital has a capacity of 150 beds and serves as the district referral hospital for a population of over 500,000 people^10^. It offers inpatient and outpatient services to the people who live in its catchment area, as well as acting as a referral center for lower-level health facilities in the district where it is located.

### Sample size and sampling technique

Using the formula N=10k/p, the sample size of the number of records to be reviewed was determined based on the event per variable (EPV) for the binary logistic regression model to establish the factors associated with patients’ misdiagnosis. In this formula, p denotes the smallest proportion of the outcome of interest, while k denotes the number of covariates (the number of independent variables) associated with the outcome. A minimum of 10 cases per variable is required for a given number of EPVs, resulting in 120 misdiagnosed cases (10 X 12) ^11^.

It has been reported that the proportion of patients who are misdiagnosed based on the opinions of an experienced physician who changes the initial diagnosis ranges between 5 and 15% ^12^. Thus, 5% was chosen as the smallest outcome proportion (p), and with 12 variables associated with misdiagnosis studied, the minimum number of patient records required was 2,400, but 31 extra records were included to account for missing data and poor record quality. Simple random sampling was used to select records from a pool of charts that met the inclusion criteria.

### Inclusion and exclusion criteria

If a patient was admitted as an inpatient through the outpatient or emergency department with a medical or pediatric non-surgical condition, their records were reviewed. A medical officer must have reviewed the patient upon admission and noted the final diagnosis, which is also considered the discharge diagnosis. This was regarded as the primary reason for the patient’s admission and treatment. There was no restriction on the age of patients’ records. Records were excluded if the patient was admitted directly to the ward (rather than through the OPD or ED) and had a surgical or obstetric condition. Furthermore, records with no evidence of a medical officer reviewing the patient, or if the patient had disappeared from the ward or been referred to a regional referral hospital, were excluded. Records of referred out patients were excluded because it was assumed that the referral to centers would possibly result into a new diagnosis, whereas for those who had died, a post mortem could be performed and possibly revealed contrasting results. Incomplete files were excluded (if they lacked more than 30% of the variables of interest), if the final discharge diagnosis was not written down by the medical officer, death while on the ward, or discharge against medical advice.

### Data collection

Four trained research assistants who were medical records officers working in two of the hospitals collected the data being supervised by the lead author. The data collectors received two days of training, which included pretesting the tools in one of the study hospitals using medical records from the 2018/2019 fiscal year. The data were collected using an explicitly designed excel-based form for some and the ODK tool for others.

#### The outcome measures

This study’s primary outcome was a patient misdiagnosis. The secondary outcomes are the factors (variables) associated with misdiagnosis obtained from previous research and experience.

#### Patient Misdiagnosis

When the admission (initial) and discharge (final) diagnoses did not match, the patient’s diagnosis status was rated as misdiagnosed. When there was concordance or matching between the admission (initial) and discharge (final) diagnoses, the diagnosis was considered correct (not misdiagnosed). In this study too, a patient was considered misdiagnosed if the admitting diagnostician prescribed treatment without a definitive or provisional (working) diagnosis.

Previous studies that investigated diagnostic error based on the difference between admission and discharge informed the choice of this definition ^14–18^. Hautz et al. (2016) investigated diagnostic discrepancy between emergency and admission departments, hypothesizing that the discrepancy was an indicator of diagnostic error^16^. This approach was used by Kijima et al (2018) to investigate the diagnostic accuracy of acutely ill elderly patients seen at a primary care center and then admitted to an emergency department within four days^14^. In a study conducted in Brazil to determine diagnostic discrepancies between admission and discharge diagnosis, Avelino-Silva and Steinman (2020) stated that, while not all of the discrepancies were due to error, a large proportion of them were^18^. Similarly, a Japanese study investigated site of infection misdiagnosis by comparing discrepancies between the initial site of infection and that assigned in the final diagnosis^15^.

During the record review, both the admission and discharge diagnoses were recorded as written on the records by the research assistant. Later, using the ICD-11 software, two of the authors, both senior clinicians, assigned the alphanumerical International Classification of Diseases Eleventh Version (ICD-11) codes. The major diagnostic grouping, parent grouping, and individual disease condition digit numbers were recorded for each diagnosis. When there was ambiguity about the codes, the two authors discussed and consulted widely until a compromise on which code to assign was reached.

### The secondary outcomes

Based on previous literature, data were extracted on variables thought to be associated with misdiagnosis^19–26^. These factors were classified as patient, disease or case-related, contextual, and health-system-related. The extracted patient data included age, gender, and whether the patient was new or old. A new patient was one who visited the health facility for the first time within one month of the admission under review, whereas an old patient was one who had visited the same hospital for either outpatient or inpatient care within one month of the admission.

The number of presenting signs and symptoms noted at the OPD/ED, the total number of final diagnoses made by the reviewing physician, and the type of illness the patient was finally diagnosed with were among the disease-related variables extracted. The diseases were classified as common and uncommon admission diseases. Common diseases of admission accounted for 20% of the conditions or diseases responsible for 80% of admissions. Malaria, Pneumonia, Severe Anaemia, Peptic Ulcer Disease, Gastroenteritis, Bacteraemia, Urinary Tract Infection (UTI), Septicaemia, Hypertension, Enteric Fever, Sickle Cell Disease with Crisis, Diabetes Mellitus, and Acute Malnutrition were among them.

All final diagnoses were arranged in the order of frequency of hospital admission, and the Pareto analysis was used to determine the diseases that accounted for 80% of the admissions. These are the diseases that were considered common admission diseases, while the rest were considered uncommon admission diseases. We also determined and gathered information on whether the initial diagnosis was made following any type of imaging and/or laboratory investigations. We also established if the patient had an underlying disease or condition, as reported by the diagnostician at the ED/OPD.

We also extracted variables related to the time of admission, such as the patient’s admission time and day of the week. The day was divided into three parts: day time (8:00 a.m. to 2:59 p.m.), evening time (3:00 p.m. to 10:49 p.m.), and night time (11:00 pm to 7:59am). The admission day was classified as either weekday (Monday through Friday) or weekend (Saturday and Sunday). The referral status of the patient, with two classifications of a referred or non-referred patient, was one of the health system-related factors. A referred patient was one who had received treatment from a lower-level private or public health facility but was later transferred to the hospital for further management, whereas a non-referred patient had not been transferred from any health facility to the hospital.

#### Data analysis

StataIC version 15.0 was used to analyze the data. Frequencies, proportions, and confidence intervals (CIs) for descriptive characteristics were reported. At bivariate analysis, the Chi square test was used to determine factors associated with misdiagnosis. A model that determined the factors associated with patient misdiagnosis was fitted using multivariable regression analysis adjusting for confounder. The level of significance was determined to be p 0.05.

### Ethical approval

This study was approved by the Department of Health Policy, Planning and Management (HPPM) of the University of Ghana (UG) with reference number HP/AC.12/1/2017. It was then approved by the Mildmay Uganda Research Ethics Committee (MUREC) (reference number is # REC REF 0505 2020) and the Uganda National Council of Science and Technology (reference number HS826ES). These were presented to the medical superintendents of the particular hospitals and the District Health Officers (DHOs) of the districts where the hospitals are located to seek for permission to conduct the study.

We applied to MUREC for waiver of patient consent to review their records since it was deemed that given the study objectives, the patients would not be harmed by accessing their information without their knowledge. Besides, we explained that since this was a retrospective paper-based records review, it would be difficult to reach out to the patients to seek for their consent since they did not provide information to the hospital that could be used to access them.

To ensure confidentiality and anonymity of the patients’ records, during the data extraction from the records, patients’ information such as their names, occupation and address were not extracted so that they could not be identified. For the purpose of easy retrieval of the variables of interest and to make it possible to crosscheck with the records, the patient’s admission number was recorded. All the data collection was conducted in the hospital in the records department and no photograph of a patient’s record was taken. No patient’s records were taken out of the hospital, and neither was any photograph of a patient’s records taken. All the study procedures were done in compliance with the Declaration of Helsinki, UNCST, MUREC and the Department of Health Policy, Planning and Management of the University of Ghana.

## Results

### Characteristics of patient records reviewed

From the 12345 records retrieved, 5832 met the inclusion criteria. Simple random sampling was used to obtain 2431 records, which were then analyzed. Up to 41.3% (n=1003) of all records (n=1003) were for children aged 0 to 9 years. Female patients outnumbered males by a slim margin (51.9% to 48.1%). The vast majority of patients (96.3%, n=2341) were admitted as new cases within the previous month. Prior to admission, the majority of patients’ diagnoses (84%, n=2041) had been made after some investigation. Up to 13% of the patients (n=317) had more than three presenting complaints. The majority of patients (84.5%, n=2055) reported no known underlying medical condition at the time of the initial diagnosis. The majority of patients (81.9%, n=1991) had one disease diagnosis recorded as the final (discharge) diagnosis while 1861 of the patients (76.6%) had a final (discharge) diagnosis(es) classified as common diseases of admission. As many as 2076 (85.4%) of the patients and 81.4% (n=1976) were admitted during the day and on weekdays (Monday through Friday) respectively. Up to 5.4% (n=131) of the patients had been referred from lower-level health facilities (Table 1).

**Table 1:** Characteristics of study variables.

| Variable |  | Frequency (f) | Percentage (%) |
| --- | --- | --- | --- |
| Hospital (N=2431) | Kiboga | 481 | 19.8 |
|  | Nakaseke | 489 | 20.1 |
|  | Mityana | 485 | 20.0 |
|  | Gombe | 489 | 20.1 |
|  | Kayunga | 487 | 20.0 |
| Age category (N=2431) | 0-9 | 1003 | 41.3 |
|  | 10 to 19 | 408 | 16.8 |
|  | 20-29 | 315 | 13.0 |
|  | 30-39 | 199 | 8.2 |
|  | 40-49 | 141 | 5.8 |
|  | Above 50 | 365 | 15 |
| Gender | Female | 1262 | 51.9 |
|  | Male | 1169 | 48.1 |
| Referral status of the patient | Referred | 131 | 5.4 |
|  | Not referred | 2,300 | 94.6 |
| Type of patient | Old patient | 90 | 3.7 |
|  | New patient | 2341 | 96.3 |
| Had initial diagnosis been made after investigations | No | 390 | 16.0 |
|  | Yes | 2041 | 84.0 |
| Number of presenting complaints | More than 3 | 317 | 13.0 |
|  | 1-3 | 2,114 | 87.0 |
| Did patient have an underlying health condition | Yes | 376 | 15.5 |
|  | No | 2055 | 84.5 |
| The number of morbidities | One disease | 1991 | 81.9 |
|  | Comorbidity (2 final diagnoses) | 390 | 16.0 |
|  | Multimorbidity (3 or more final diagnoses) | 50 | 2.1 |
| Time of admission | Day time | 2,076 | 85.4 |
|  | Evening | 223 | 9.2 |
|  | Night | 132 | 5.4 |
| Period of the week patient was admitted | Weekday | 1976 | 81.3 |
|  | Weekend | 455 | 18.7 |
| How common is the diagnosis made | Combination of common and uncommon disease | 144 | 5.9 |
|  | Only common disease | 1861 | 76.6 |
|  | Only uncommon disease | 426 | 17.5 |
| Diagnostic status | Not misdiagnosed | 2208 | 90.8 |
|  | Misdiagnosed | 223 | 9.2 |

### Outcomes

Overall, 223 (9.2%; 95% CI: 8.1-10.3%) were misdiagnosed. Nakaseke hospital accounting to 27.8% (62) of all the 223 misdiagnosed cases while Kiboga hospital had the lowest (14.4%) of the misdiagnosed cases (Table 2).

**Table 2:**
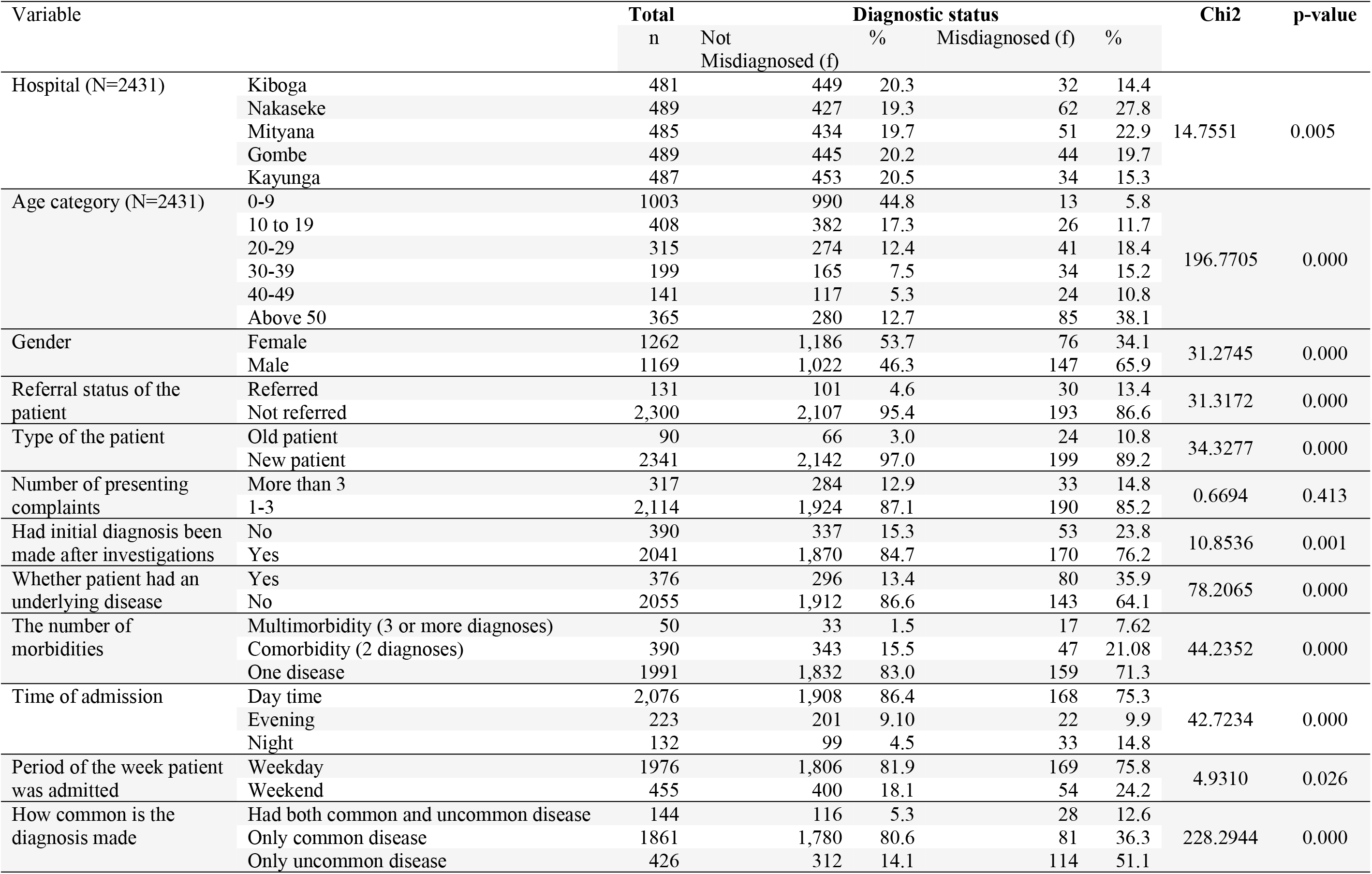
Cross-tabulation of independent variables and misdiagnosis of in-patients based on the Chi-square test statistic with corresponding p-value in general hospitals of Central Uganda.

### Factors associated with misdiagnosis

At bivariate analysis, misdiagnosis was associated with hospital (p=0.005), age (p0.001), gender (p<0.001), referral status of the patient (p<0.001), patient type (p<0.001), and whether initial diagnosis was made after some investigations (p=0.001). Other factors included whether the patient had an underlying disease (p<0.001), the number of morbidities (p<0.001), the time of admission (p0.001), the period of the week the patient was admitted (p=0.026), and the frequency with which the diagnosis was made (p<0.001).

Factors associated with misdiagnosis in multivariable analysis included the hospital of admission, the patient’s age, gender, referral status, number of morbidities, time of day when the patient is admitted, and the type of disease (es) the patient had as the final diagnosis (Table 3).

**Table 3.** Multivariate logistic regression on factors associated with inpatients misdiagnosis in general hospitals of Central Uganda (N=2431)

| Variable |  | Misdiagnosed<br>n | % | Odds<br>Ratio | P value | 95% CI |  |
| --- | --- | --- | --- | --- | --- | --- | --- |
| Hospital | Kiboga | 32 | 14.4 | Reference |  |  |  |
|  | Nakaseke | 62 | 27.8 | 1.95 | 0.01 | 1.17 | 3.25* |
|  | Mityana | 51 | 22.9 | 1.40 | 0.213 | 0.83 | 2.37 |
|  | Gombe | 44 | 19.7 | 0.98 | 0.956 | 0.57 | 1.69 |
|  | Kayunga | 34 | 15.3 | 0.90 | 0.722 | 0.51 | 1.59 |
| Age | 0-9 | 13 | 5.8 | Reference |  |  |  |
|  | 10 to 19 | 26 | 11.7 | 4.61 | 0.0000 | 2.30 | 9.25* |
|  | 20-29 | 41 | 18.4 | 8.15 | 0.0000 | 4.18 | 15.89* |
|  | 30-39 | 34 | 15.2 | 8.12 | 0.0000 | 3.99 | 16.54* |
|  | 40-49 | 24 | 10.8 | 7.88 | 0.0000 | 3.71 | 16.73* |
|  | 50+ | 85 | 38.1 | 12.14 | 0.0000 | 6.41 | 23.01* |
| Sex | Female | 76 | 34.1 | Reference |  |  |  |
|  | Male | 147 | 65.9 | 1.89 | 0.0000 | 1.35 | 2.64* |
| Referral status | Referred | 30 | 13.4 | Reference |  |  |  |
|  | Not referred | 193 | 86.6 | 0.51 | 0.011 | 0.31 | 0.86* |
| Case type | Old patient | 24 | 10.8 | Reference |  |  |  |
|  | New patient | 199 | 89.2 | 0.94 | 0.85 | 0.51 | 1.75 |
| Lab tests done | No | 53 | 23.8 | Reference |  |  |  |
|  | Yes | 170 | 76.2 | 1.18 | 0.41 | 0.79 | 1.76 |
| Underlying disease | Yes | 80 | 35.9 | Reference |  |  |  |
|  | No | 143 | 64.1 | 0.63 | 0.015 | 0.43 | 0.91* |
| Number of morbidities | One disease | 17 | 7.62 | Reference |  |  |  |
|  | Comorbidity (2 diseases) | 47 | 21.08 | 1.49 | 0.1 | 0.93 | 2.38 |
|  | Multimorbidity | 159 | 71.3 | 4.71 | 0.001 | 1.91 | 11.65* |
| Time of admission | Day time | 168 | 75.3 | Reference |  |  |  |
|  | Evening | 22 | 9.9 | 1.26 | 0.10 | 0.74 | 2.14 |
|  | Night | 33 | 14.8 | 3.00 | 0.000 | 1.81 | 5.02* |
| Period of the week | Weekday | 169 | 75.8 | Reference |  |  |  |
|  | Weekend | 54 | 24.2 | 1.23 | 0.279 | 0.84 | 1.79 |
| How common the diagnosis is | Combination of common and uncommon disease | 28 | 12.6 | Reference |  |  |  |
|  | Only common disease | 81 | 36.3 | 0.55 | 0.083 | 0.28 | 1.08 |
|  | Only uncommon diseases | 114 | 51.1 | 2.57 | 0.008 | 1.28 | 5.18* |

After controlling for other variables, the results revealed that the odds of being misdiagnosed were two times higher in patients admitted to Nakaseke Hospital than in patients admitted to Kiboga Hospital (adjusted Odds (aOR) = 1.95, 95% CI = 1.17-3.25, p=0.01When compared to being admitted during the day, being admitted at night was associated with an increased likelihood of being misdiagnosed (aOR 3.0, 95% CI: 1.81-5.02, p0.01). Male patients had an 89% higher chance of being misdiagnosed (aOR=1.89 95% CI=1.35-2.64, p0.01) than female patients. Patients aged 10-19, 20-29, 30-39, and 40-49 were more likely to be misdiagnosed 2.3 times (aOR=2.3, 95% CI: 2.3-9.25, p0.01), 8.2 times (aOR=8.2 95% CI=4.18-15.89, p0.01), 8.12 times (aOR=8.12 95% CI=3.99-16.54, p0.01), 7.88 times (aOR=7.88 95%.

Having no underlying diseases was associated with a 37% lower risk of misdiagnosis (aOR=0.63; 95% CI: 0.43-0.91, p=0.015). When compared to having a single morbidity, having multiple morbidities was associated with a 5 times greater likelihood of being misdiagnosed (aOR=4.71; 95% CI: 1.91-11.65, p0.01). Patients treated for uncommon diseases of hospital admission were approximately three times more likely to be associated with misdiagnosis than patients treated for both common and uncommon diseases of admission (aOR=2.57; 95% CI: 1.28-5.18, p0.01). When compared to patients referred from other lower-level health facilities or clinics, patients who had not been referred had a 49% lower likelihood of misdiagnosis (aOR=0.51; 95% CI=0.31-0.86, p=0.011).

### Diseases/conditions involved in misdiagnosis

Table 4 shows the ICD-11 codes of the initial (misdiagnosed) and final (correct) conditions or diseases.

**Table 4:** The ICD-11 coding of the initial and final diagnoses for the misdiagnosed patients (n=223)

| Initial (wrong) diagnosis | Final (right diagnosis) | Initial (wrong) diagnosis | Final (right diagnosis) |
| --- | --- | --- | --- |
| 1. QA1C | MG24.1 | 111.DA42.Z; MB24.3 | 1A40.Z |
| 2. MA15.0; FA20.Z | 8B20 | 112.MG40.Z | MG45.Z |
| 3. 1F40.Y; 1A07.Z; MA15.0; DA61 | GB51 | 113.CA40.Z; 1B70.Z; FA05 | CA22.Z; BD10&XT5R; 8A61.3Y |
| 4. 1F40.Y; DA42.Z | 1A07.Z; DA61; 1A40.Z | 114.FA8Z; CA45; MF50.2Y | FA80.9; CA20.Z; 5B81.01 |
| 5. QA1C | 1B95 | 115.QA1C | MB21.4; 3A9Z |
| 6. 1A90; AA8Z XT5R | 1E91.Z; 1C62.3; 1F2Z | 116.CA20.Z; GCO8.Z | 1B10.1; 3A9Z; 5B54 |
| 7. IC62.Z | GA90 | 117.BC4Z; DA61 | BD10 |
| 8. 1A40.Z | 1A1Z | 118.GB01.1 | GC00.Z; GB02.Z |
| 9. CA23 | CA22.0 | 119.QA1C | 8A80.Z |
| 10. QA1C | 1F40.Z | 120.GB41; BD10 | GB40; 9C80.0 |
| 11. QA1C | 5A41; 6A70.Z | 121.1B98; DA61; DA42.Z; 4A84.Z | 1F57.Z |
| 12. GA05.Z; GC00.Z | GCO8.Z | 122.QA1C | 5B7Z; 1B10.0; 5C70.0 |
| 13. CA40.07 | 1B10.0 | 123.QA1C | CA40.Z&XT6S&XY69 |
| 14. PA78&XE11V | 1B70.Z | 124.MD81.3; ME24.90; 1B12.7 | 1A40.Z; DC50.0 |
| 15. QA1C | DC12.Z&XT8W | 125.QA1C | 1E50.1; DA61 |
| 16. QA1C | 1A40.Z | 126.QA1C | 8B20; BA00.Z |
| 17. BD11.0 | BD13 | 127.GCO8.Z; GB51 | GC00.Z |
| 18. QA1C | 1F40.Y | 128.1A40.0 | ME05.1 |
| 19. QA1C | 1F40.Y | 129.AB0Z | AA91.Z |
| 20. QA1C | 5A41 | 130.QA1C | MA15.0 |
| 21. QA1C | 1F40.Y | 131.3A9Z | 8B20; 8A6Z |
| 22. 3A9Z | DA61 | 132.BD1Z | 5DOY; BA00.Z |
| 23. DA61; 1F40.Y | 1A07.Z; 1B95; CA45 | 133.6C40.3; 5C70.0 | 5B5A.10; 5C70.0; CA40.Z |
| 24. QA1C | DA61 | 134.GCO8.Z | 1F40.Y |
| 25. QA1C | MA15.0 | 135.QA1C | 1F45; GC00.Z |
| 26. 1F45; GCO8.Z | 1B10.1 | 136.QA1C | DA61 |
| 27. QA1C | CA40.Z&XT6S&XY69 | 137.1F45 | GCO8.Z |
| 28. 1F40.Y; 1A07.Z; 1E50.Z | GB51 | 138.1F45 | 1A40.Z |
| 29. 6B01; 6C20.Z | 6B60.Z | 139.1F40.Y | 1A40.Z; BA00.Z |
| 30. NE60 | EB13.0 | 140.QA1C | 1F45 |
| 31. GB60.Z; 4B40; 2C94.Z | GB61.Z | 141.CA22.Z | DA61 |
| 32. ME84.2Z; DA61; NB9Z | FA8Z | 142.QA1C | 1A07.Z; 3A9Z |
| 33. BC20.1; BB01.Z | BD11.Z | 143.1B10.Z | CA40.Z |
| 34. 1C62.Z | 1C62.3 | 144.1A1Z | DA61 |
| 35. 1F45; 1C41; 1A36.00; 1A07.Z | BA00.Z | 145.DA42.7 | 1F45; DA61 |
| 36. 1A07.Z | DC50.Z | 146.6C40.3; 3A9Z | NA0Z; 6C40.3 |
| 37. 1B10.1 | CA40.Z&XT6S&XY69 | 147.QA1C | 1F40.Y |
| 38. 1A40.0 | 1A07.Z | 148.QA1C | 1F40.Y |
| 39. QA1C | 1F40.Z | 149.CA45 | 1B10.1 |
| 40. GA05.Z | DA61 | 150.3A9Z | 5A21.Z; BA00.Z |
| 41. MA15.0 | DA61 | 151.8C01.3; FA92.Z | FA2Z; GCO8.Z |
| 42. 1F45; 5A41 | 6A20.0Z; 8A6Z | 152.CA40.Z | GC00.Z; 3A9Z; ME05.0 |
| 43. QA1C | BA00.Z | 153.BA00.Z; 8C0Z | BA01 |
| 44. DA61 | 1B95 | 154.6A2Z; 5A14 | BA03; 5A14 |
| 45. 1A40.Z; DA61 | DA42.Z | 155.BD10; CA20.Z | BA6Z |
| 46. QA1C | DA61 | 156.GB6Z; DA61; 5B5A | 1A40.Z; 1A07.Z; DB94.Z; MA18.0 |
| 47. 6C4Z | 6C4G.70; BA00.Z | 157.DA42.Z | DA61; 1A07.Z |
| 48. QA1C | 6D10.Z | 158.1F45; MA15.0 | 1B10.1; 1F23.0 |
| 49. QA1C | 5A14 | 159.CA45 | GB6Z; GB61.Z; DA61 |
| 50. 8A68.Z | 8A6Z | 160.6C2Z | 1A40.Z; 6A2Z |
| 51. QA1C | 1F45 | 161.QA1C | 1A40.Z; CA09.Z |
| 52. QA1C | GCO8.Z; FA92.Z | 162.QA1C | ME05.1 |
| 53. 5A14; BA00.Z; BC4Z | 5A21.0 | 163.QA1C | CA40.Z&XT65 |
| 54. 4A84.Z | CA71.0 | 164.1B40.Z | 1A07.Z |
| 55. 2C13.0 | DD71.Z | 165.DA61 | 6A60.9 |
| 56. 1F40; CA71.0; 5A22.Z | CA40.Z&XT65 | 166.1F40.Y; MA15.0 | FA8Z |
| 57. DA61; BA00.Z; 3A9Z | GB5Y | 167.1F40.Y | MA15.Y |
| 58. BA00.Z; 1F45 | BD90.Y | 168.8B20; MA15.0 | 5A41 |
| 59. CA40.Z | 8C03.0; BA00.Z | 169.DA61 | 1C41 |
| 60. 1F40 | 1A07.Z | 170.1B10.1; BD10 | BA01; CA07.0 |
| 61. CA40.Z | CA22.Z; 1B70.Z | 171.5C70.0; 1B10.1 | CA45 |
| 62. QA1C | 1A07.Z | 172.5C70.0; 1C41 | 1A40.Z |
| 63. CA45; 5C70.0 | 1A07.Z; CA40.Z; 1A40.Z | 173.CA40.Z; CB03.4 | MD11.6 |
| 64. BD10; GC2Z; DB98.7Z | BD13; BB01.5 | 174.MC81.2 | 6D71; BD11.Z |
| 65. 1D01.10 | GCO8.Z; 8B42; 1A07.Z | 175.DA61 | DA42.Z; 1E90.0 |
| 66. 5B7Z; 5C70.0 | 5B52&XS25; 1A40.Z | 176.CA40.Z; 1C60.Z | 1C62.3; 3A9Z |
| 67. QA1C | 1A40.Z | 177.BD10 | MA15.Y |
| 68. 5A11 | DA42.Z; GCO8.Z; 1F45 | 178.CA07.0; 1F23.0 | CA40.Z; DA61 |
| 69. MB24.5 | 1A62.0; 2B31.20 | 179.CA07.0 | BA00.Z; CA40.Z; GCO8.Z; 1A07.Z |
| 70. CA23.30 | CB03.Z | 180.ME05.1 | 1A40.Z; DA61 |
| 71. MG45.Z; 1F45; GCO8.Z | DA22.Z; DA61 | 181.6D85.3; 1B10.1 | 1D01.10; CA07.0 |
| 72. BA00.Z | BE2Y | 182.DA61; 1A07.Z; DB94.3 | DB94.3; 3A9Z; GCO8.Z |
| 73. 6C40.Z; 5A14 | 1C8E; 5A21.Z | 183.CA40.Z; 1B10.1 | CA07.0; 6A60.9 |
| 74. CA20.1Z; 8B20 | CA22.Z | 184.5A11; DA42.Z | GCO8.Z; DA61 |
| 75. MA15.Y; 1B10.1 | DA42.82 | 185.5A11 | DA61 |
| 76. MA15.Y | BD10 | 186.CA07.0; 5C70.0 | CA40.Z |
| 77. QA1C | GCO8.Z | 187.1A07.Z | MA15.0 |
| 78. 1C41 | 8A80.Z; MA15.Y | 188.DA61 | DC50.Z |
| 79. 3A9Z | MD31; DA61 | 189.QA1C | 1F40.0 |
| 80. DA42.Z; CA07.0; 1B10.1 | PD03 | 190.QA1C | 1B10.1 |
| 81. QA1C | QA14 | 191.1A40.Z; MA15.0 | ME24.9Z |
| 82. 4B23; 1B10.1; GCO8.Z | CA40.Z | 192.6A20.Z | 9C83.13 |
| 83. CA40.Z | BD10 | 193.MA15.0 | MA15.Y |
| 84. CA45; 5C70.0 | MA15.Y | 194.BA00.Z | BD10 |
| 85. 5C70.0; 5B52&XS25 | DA22.Z | 195.GC00.Z | ME10.02 |
| 86. MD81.3 | DA61 | 196.QA1C | 2C77.Z |
| 87. QA1C | MA15.Y | 197.CA23.30 | 1B10.1 |
| 88. 1C41; DA42.Z | CA45 | 198.BA00.Z; DA61; CA40.Z; CB27 | BA01; 5A11; CA22.Z |
| 89. MA15.0 | MA15.Y | 199.MA15.0 | 5A11 |
| 90. QA1C | MG22 | 200.DA61 | BA03 |
| 91. 1C62.Z; 1F23.2; CA07.0 | CA45; 1D01.10, 1B10.1 | 201.MG40.Z | 1F40.0 |
| 92. DA61; DB93.1 | DB95.Z | 202.MF50.3; 1A6Z | BE2Y |
| 93. CA07.0 | MA15.0 | 203.BE2Y; 1F40; CA40.Z | 1F57.Z |
| 94. 3A51.2 | 1F40.0; GCO8.Z | 204.MA15.0 | 1D01.10 |
| 95. KA60 | KB8Z | 205.5A11 | MA15.0 |
| 96. MA15.Y | KA87.Y | 206.5A11 | DA42.Z |
| 97. MA15.Y | KB06 | 207.8A81.Z | 6A7Z |
| 98. DA61; BA2ZZ; 3A9Z&XS25 | 2B33.1 | 208.CA07.0 | 5B7Z |
| 99. GB40 | GB61.Z | 209.GB40 | BD10 |
| 100.QA1C | CA40.Z | 210.1C62.Z | 6D85.3 |
| 101.DA61; 1C41; CA07.0; GCO8.Z | 8B42 | 211.5C70.0 | 8B20 |
| 102.MC81.2 | BD10 | 212.QA1C | 1F45 |
| 103.QA1C | CA45 | 213.1F40.Y | 5C70.0; 3A9Z |
| 104.ME05.1; 5B7Z | GB61.Z; 1A40.Z | 214.1A07.Z | 1E50.Z |
| 105.ME04.Z | BD10 | 215.GB02.Z; GCO8.Z | 1A6Z |
| 106.1F40.Y | GB51 | 216.CA00 | CA08.1Z |
| 107.5A11 | 5B57.Z | 217.QA1C | MA15.0 |
| 108.MA15.0 | 1D01.10 | 218.GB40 | GB61.Z |
| 109.QA1C | 1A40.Z | 219.QA1C | CA40.Z |
| 110.1A07.Z | GA34.3 | 220.DA61; 1C41; CA07.0; GCO8.Z | 8B42 |
| 111.3A9Z&XS25 | 3B4Z | 221.MC81.2 | BD10 |
| 112.MA15.0 | CA22.Z | 222.MC81.2 | 6D71; BD11.Z |
| 113.QA1C | 1F23.0 | 223.4A84.Z | CA71.0 |

## Discussion

The current study sought to determine the magnitude of patient misdiagnosis as well as the relationship between patient, disease, and health-care system factors associated with patient misdiagnosis in general hospitals in central Uganda. According to the findings, 9.2% of inpatients were misdiagnosed at the outpatient and emergency departments, which are the points of admission. Being a patient in the older age bracket and being a male patient were the patient-related factors associated with misdiagnosis. Misdiagnosis was associated with disease-related factors such as being treated for an uncommon condition, having an underlying disease, and having multimorbidity (three or more diseases). Being admitted at night was one of the environmental factors associated with patient misdiagnosis. A patient being referred was one of the health-care system factors associated with misdiagnosis.

The findings of this study revealed that older patients were more likely to be misdiagnosed than younger patients. This finding is consistent with the belief that there have been deep flaws in health-care systems in the delivery of health-care services to older patients ^27^. Older patients may be more complex and difficult to diagnose and treat than younger patients hence the increased chances of misdiagnosis ^28^. It has been reported, for example, that older patients typically have close to five medical conditions and possible failures, which may weaken other body systems ^29^.

The better diagnostic outcome performance in children aged 0-9 years compared to patients of other ages brackets is most likely explained by increased efforts in improving the management of Uganda’s high morbidity burden childhood illnesses such as malaria, pneumonia, diarrhoea, and anaemia ^30^. This has included the distribution of rapid diagnostic testing kits, the training of frontline health workers in the management of these diseases, and the improvement of referral systems for such illnesses.

The study also discovered that male patients were roughly twice as likely as female patients to be misdiagnosed. This finding is somewhat surprising given that previous research, which was mostly descriptive in nature, hypothesized that even when they presented with similar symptoms, females were more likely to be misdiagnosed than males ^8,31,32^. Nonetheless, there are a number of possible explanations for this discovery. Because most health-care systems have traditionally prioritized women’s and children’s health, misdiagnosis in men and the elderly may be inescapable because health-care workers are less familiar with their problems^33^. Another possible explanation for this finding is that men, in comparison to women, are less involved in the diagnostic decision-making process due to their lack of experience with healthcare systems. Their interactions with diagnosticians will eventually become less eventful, resulting in poor diagnostic outcomes ^34^. This suggests that in order to achieve better diagnostic outcomes, men’s health should be improved, as well as their health-seeking behaviors and participation in the diagnostic decision-making process. According to the findings, patients with multimorbidity were approximately 5 times more likely to be misdiagnosed. The findings of this study are consistent with those of Panagioti et al. (2015), who discovered that patients with multimorbidity were 1.2 times more likely to have a diagnostic error ^35^. According to Aoki and Watanuki (2020), while misdiagnosis occurred in 3.9% of patients without multimorbidity, it occurred in 4.9% of multimorbid patients in Japan ^36^. Diagnostic decision-making in multimorbid patients can be difficult if clinicians are unfamiliar with the cases that multimorbid patients may present with, as observed in this study. Clinicians’ attention may be drawn to the most obvious complaints, especially if the patient has an underlying disease, and they may miss any subtle signs.

In terms of underlying diseases and their relationship to misdiagnosis, this study discovered that patients with underlying diseases were more likely to be misdiagnosed than patients who did not report having any. The presence of underlying diseases raises the possibility of bias(es) in diagnostic decision making. In one sense, if a patient mentions an underlying disease, diagnosticians may follow that line of thought and disregard thinking outside the box of the underlying disease. On the other hand, if the underlying disease is not mentioned, clinicians may not think along those lines, despite the fact that it is the most likely pathology. Hausmann et al. (2019) alluded to this possibility as one that could lead to ambiguity and misdiagnosis^37^.

The night shift was found to be the most common time of day for patient misdiagnosis. This finding broadly supports the findings of other studies in this field, which have linked the night shift to the occurrence of diagnostic errors ^38,39^. For example, Hughes (2015) reported that error rates among nurses were more likely to occur at night than during the day, citing sleepiness, anxiety, and attention deficit as possible causes of error^39^. Health workers are frequently fatigued at night, resulting in poor cognitive performance^40^. Similarly, sleep deprivation has been identified as a leading predictor of errors committed during night shifts ^41^.

There was no significant association found in this study between misdiagnosis and patients being admitted on weekends or weekdays. Weekend shifts have previously faced challenges such as insufficient staffing and logistics^42^. This could imply that misdiagnosis was unaffected by the day of the week in the hospitals studied. This explains why vigilance for quality diagnosis and measures for better diagnostic practices should be maintained at all times. Previous research has also shown that patients who present on the weekend are more likely to be critically ill^43^. If this had occurred in the hospitals studied, such patients would have been referred to other higher-level hospitals for further care.

Patients who were not referred from lower-level primary healthcare facilities had a lower risk of misdiagnosis. The primary reason for the likely association with misdiagnosis for referred patients is a cognitive bias in which receiving clinicians were predisposed to accept the referring diagnostician’s diagnosis^21^. If the referring diagnostician was incorrect and the receiving diagnostician continues to use the same diagnosis without further inquiry, this results in a misdiagnosis. Furthermore, a study of diagnostic errors in tuberculosis patients found that misdiagnosis was common among referred patients due to uncoordinated referral systems in which the referring physician provided little information about the reasons for referral to the receiving physicians^44^. Furthermore, referral cases are more likely to be complicated, making diagnosis more difficult to negotiate than non-referred cases^45^. By paying special attention to referred cases during the diagnostic process, the quality of diagnosis in this group of patients is likely to improve.

### Implication of the findings and future direction

The study’s findings could be used by hospital administrators to ensure that quality diagnostic standards are followed at all times during hospital operations, including weekends and nights. Hospitals should prioritize identifying bottlenecks to quality diagnostic decision making using a system approach. Hospital administrators should ensure that hospitals remain safe places for patients by doing everything possible to ensure appropriate diagnostic outcomes. As part of the key quality improvement parameters, they should perform their supervisory and stewardship roles and make diagnostic errors. They should also make it a part of the hospital’s patient safety culture. These findings should be used for learning purposes by diagnosticians in hospitals and other levels of the health care system, particularly on the factors that predispose one to error. To ensure better management of patients with complex conditions, such as those with comorbidities and underlying diseases, coordinated efforts are required. It is recommended that clinicians be trained on how to anticipate, diagnose, and manage patients with whom certain factors were found to be associated with misdiagnosis in this study.

### Study limitations

This study’s findings are subject to at least three limitations. First, because the records reviewed were paper-based, there were numerous errors that resulted in the exclusion of many records. This may have influenced the findings. Second, because the study sites were limited to general hospitals, the results may not be representative of what happens at higher levels of the healthcare system, such as regional and national hospitals, where disease profiles are likely to differ. Lastly, we were unable to locate information on the diagnosticians. As a result, it would be prudent to conduct research at higher levels of the healthcare system, as well as research that takes diagnosticians’ variables into account.

## Conclusions

These findings have significant implications for patient care, including providing information that aids understanding of how misdiagnosis and diagnostic error manifest in hospitals, as well as where emphasis should be placed to improve diagnostic decision making.

## Author contributions

All authors made a significant contribution to the conception, study design, execution, acquisition of data, analysis and interpretation of the work reported. They also took part in drafting, revising or critically reviewing the article. They as well gave final approval. They agree to be accountable for all aspects of the work.

## Acknowledgments

We acknowledge the medical superintendents who gave permission for the study to go on in their hospitals. We also acknowledge the records officers and hospital staff who played a part in this study by retrieving the medical records and those who participated as data collectors.

